# In search of an honest butterfly: sexually selected wing coloration and reproductive traits from wild populations of the cabbage white butterfly

**DOI:** 10.1101/2021.03.07.434307

**Authors:** Anne E. Espeset, Matthew L. Forister

## Abstract

Sexual selection is central to many theories on mate selection and individual behavior. Relatively little is known, however, about the impacts that human-induced rapid environmental change are having on secondary sexually selected characteristics. Honest signals function as an indicator of mate quality when there are differences in nutrient acquisition and are thus potentially sensitive to anthropogenically-altered nutrient inputs. We used the cabbage white butterfly, *Pieris rapae* (L.), to investigate differences in color and testes size in a system that is often exposed to agricultural landscapes with nitrogen addition. We collected individuals from four sites in California and Nevada to investigate variation in key traits and the possibility that any relationship between wing color and a reproductive trait (testes size) could vary among locations in the focal butterfly. Coloration variables and testes size were positively albeit weakly associated across sites, consistent with the hypothesis that females could use nitrogen-based coloration in the cabbage white as an indicator for a male mating trait that has the potential to confer elevated mating success in progeny. However, variation in testes size and in the relationship between testes size and wing color suggest complexities that need exploration, including the possibility that the signal is not of equal value in all populations. Thus these results advance our understanding of complex relationships among environmental change and sexual selection in the wild.

---

Some of the most dramatic and inspiring aspects of the animal world are sexually selected traits, including mammal horns, bird feathers, and the colors of insect wings, which have all been appreciated by natural scientists and human societies (Darwin 1896, Johnstone 1995). Thus, it is not surprising that a range of theories have been developed to make sense of this diversity, with one of the most influential being Zahavi’s handicap principle of sexual selection (Zahavi 1975). This is the idea that because secondary sexual signals are costly and not all individuals are able to produce these expensive traits, their presence serves as an honest signal of individual quality to potential mates. The handicap principle of sexual selection has been successfully investigated in a number of natural systems. For example, in many bird species it has been observed that longer tails (Moller and de Lope 1994) and testosterone-dependent ornaments (Folstad and Karter 1992, Mougeot et al. 2007) are indicators of individual quality.

In addition to assuming variation among individuals in quality and the existence of expensive traits, honest-signaling theory relies on the ability of the signal receiver to distinguish high-quality signals as well the existence of a benefit for the signal receiver for mating with such individuals. In some cases, the benefit may be directly realized through nuptial gifts or other such provisioning (Arnqvist and Nilsson 2000, Stålhandske 2001, Fedorka and Mousseau 2002, Lewis et al. 2014). In other cases, the benefit may be through genes that will be passed on to offspring with associated improvements to their fitness, as in the sexy sons hypothesis (Weatherhead and Robertson 1979). For instance, individuals that can expend a higher amount of energy acquiring nutrients for a signal show potential mates that their offspring will also be able to expend similar amounts of energy to obtain costly nutrients (Scriber and Slansky 1981). The ability to acquire nutrients has impacts on individual fitness (Boggs and Freeman 2005) and population dynamics (Batzli 1983, Pedersen and Greives 2008) and may be especially important under stressful conditions including in areas exposed to high levels of human-induced change (Boggs and Freeman 2005, Boggs 2009, Boggs and Niitepõld 2014).

In the era of human-induced rapid environmental change, examples are being reported in which honest signals and associated “handicaps” have changed or might be affected in coming years (Sih et al. 2011, Tuomainen and Candolin 2011). Of particular interest in this context is the fact that human activity, through industry, agriculture and private land management, is having enormous effects on nutritional landscapes (Snell-Rood et al. 2015). In disturbed areas, such modification comes about through food waste (Robb et al. 2008), road salting (Snell-Rood et al. 2014), vehicle-associated aerial chemical deposition, and fertilizer application, among other processes. Fertilizer deposition and pollution are flooding natural and disturbed areas with once-limiting nutrients, such as phosphorous and nitrogen (Vitousek et al. 1997, Smil 2000, Galloway et al. 2008, Yuan et al. 2018), and our understanding of the implications of such nutritional changes on sexually selected characteristics is growing (Snell-Rood et al. 2015), especially in terms of honest signaling and sexual selection (Goos et al. 2016, Espeset et al. 2019).

In this study, we ask if a putative honest signal is associated with a measure of potential mating success (testes size, described further below), and if that relationship (between advertisement and mating potential) differs among populations. *Pieris rapae* (L.), the cabbage white butterfly, was chosen as our model system because of its mate-choice strategy and previous information on sexually selected characteristics of this species. For example, we know that *P. rapae* females select mates based on wing coloration (Morehouse and Rutowski 2010b) and that the characteristic white and yellow wing color of butterflies in the family Pieridae is from the pterin (or pteridine) pigment compound class (Stavenga et al. 2004, Wijnen et al. 2007). Pterins are made up of heterocyclic compounds packed with nitrogen; thus, all individuals who have high amounts of pterin pigments in their wings must have had access to sufficient amounts of nitrogen. We can expect that the availability of sufficient nitrogen would mean that organisms could invest in advertising (secondary sexual traits) while still investing in basal reproductive traits, such as testes size, thus producing a correlation among individuals between secondary sexually-selected traits and reproductive value or potential.

In this context, we consider testes size to be an indicator of reproductive potential because males with larger tests are more likely to sire offspring, either through enhanced sperm competition or elevated mating frequency (Vahed and Parker 2012). Thus females that choose males with larger testes increase their chances of having sons that mate successfully, increasing their own fitness, and this is expected to be especially true in more polyandrous butterflies (Gage 1994) which includes *P. rapae* (Bissoondath and Wiklund 1996). It is important to note that this picture of honest signaling and sexual selection may shift if all individuals in a population have access to excessive amounts of a previously limiting nutrient (discussed further below).

In this study we address the following: (1) Does cabbage white wing coloration predict testes size? This is a key element of honest signaling theory applied to the cabbage white system, and addressing this question is our first motivation for the present study. Secondarily, we ask: (2) Do wing coloration, sexual organs, and the relationship between coloration and a reproductive trait vary among populations located near high agricultural areas and areas with lower agricultural activity? (3) Finally, we investigate variation in spermatophore count as an index of repeated matings among study sites which potentially provides information on variation in mating behavior.

In general (and in answer to question 1), we hypothesize that butterflies with larger testes will have increased brightness, hue, and saturation, signifying the honesty of wing coloration for reproductive quality as a trait that females could pass on to their sons and thus increase their evolutionary fitness. Additionally (in answer to question 2), we could expect to see butterflies with larger testes and more colorful wings to be from populations with higher atmospheric nitrogen deposition (e.g. developed and highly populated areas) or land use associated with a higher direct fertilizer input (e.g. croplands). Similarly, we anticipated seeing less variation among individuals from high-nitrogen areas and greater variation among individuals from natural or undisturbed areas due to the more heterogeneous nature of nutrient availability in non-agricultural areas (see Fig. 1 for an illustration of one possibility in which these among-population differences might be realized). Furthermore, if individuals from high-nitrogen areas indeed have uniformly larger testes and brighter wings, then it is possible that the honest signaling relationship between testes and coloration will be absent in these areas (Fig. 1B), and even (finally) it is possible that sexual selection will thereby have been relaxed on wing coloration and individuals in high-nitrogen areas could actually be less bright relative to natural areas, if selection on wing color has been relaxed (the latter possibility is not illustrated in Fig. 1, see note in legend). We raise these possibilities while acknowledging that other factors (besides anthropogenic nitrogen addition) are likely important, including host plant availability and climatic differences. An investigation of these factors would require a great many more locations than are studied here. Thus we present the agricultural setting of our populations as a context in which to view the results and do not aim for a direct hypothesis test of the effect of agriculture on reproductive traits (either primary or secondary).

**Fig. 1.**
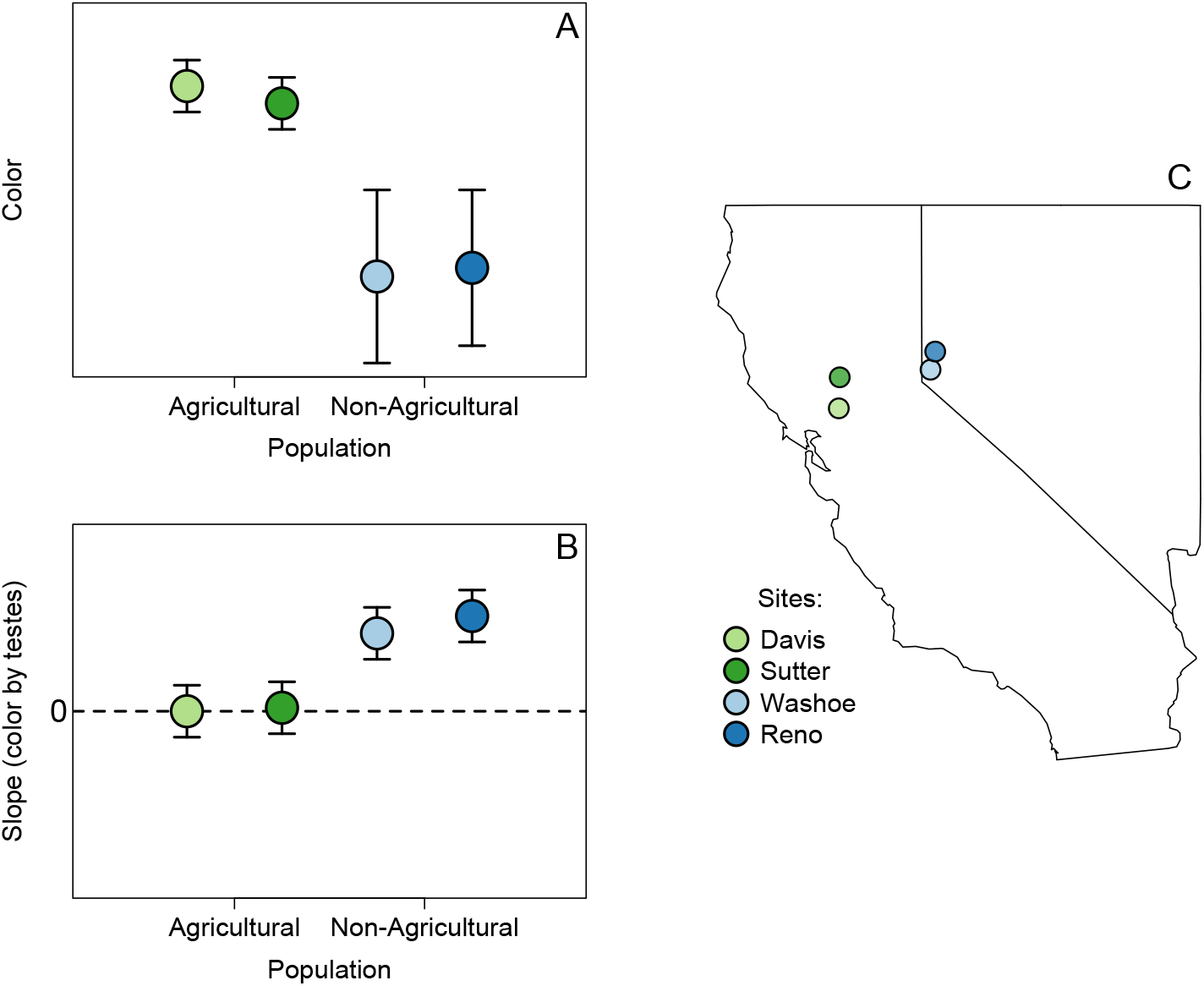
Expectations for the evolution of an honest signal (A and B) and a map (C) of collection locales. Values graphed in panels (A) and (B) are artificial, made to illustrate the following ideas. In (A), wing color by land use, we expect an increase in color in association with higher nitrogen availability at agricultural sites (green points). We also hypothesize a smaller variance within these populations because all individuals have access to excess nitrogen. In (B), slopes of the relationship between color by testes weight are a measure of signal honesty (more positive slopes indicate a relation between traits in which males with more reproductive potential are also more intensely colored). In areas where there is excess nitrogen (agricultural areas) we would expect the correlation of color by testes weight to be zero or closer to zero (because the signal has lost value). In more nitrogen-limited areas, in contrast, we would expect to see a positive relationship between color and testes weight corresponding to signal honesty. Points in (C) are actual collection locations of *P. rapae* obtained across different proportions of cropland in Northern California and Northern Nevada. Blue represents sites and areas with low proportion of cropland (Washoe and Reno) and green signifies high levels of cropland (Davis and Sutter). Note that the expectations illustrated in the first two panels are a potential snapshot of a complex process. A further possibility (not illustrated here) is that loss of the honest signal in populations where all individuals have access to excess nitrogen could ultimately relax selection on wing coloration and lead to reduced brightness in those areas (i.e., a further step in the process than illustrated in panel (A).

## Materials and Methods

### Specimens

Butterflies were collected at two sites in northern California and two in northern Nevada between June and July 2018 (Figure 1) and transported in glassine envelopes to a lab at the University of Nevada, Reno. Sites were selected to encompass variation in land use intensity, especially regional agricultural activity. With respect to surrounding land-use, our Washoe site had the lowest proportion of cropland (0.125%) within a 10km buffer around our collection location, followed by Reno (0.416%), then Davis (71.1%), and finally Sutter (85.9%) with the highest proportion of cropland (National Land Cover Database; www.mrlc.gov).

As mentioned above, our goal was not to formally test for a difference between agricultural and non-agricultural sites, rather we encompassed this landscape variation in order to have the most general picture possible (given a modest population-level sample size) of the relationships among our focal traits. Once in the lab, bodies and wings were separated; wings were stored in a dark drawer to avoid light degradation and bodies were stored in a −20°C freezer until further processing.

### Wing Measurements

Wing coloration was measured using a JAZ spectrophotometer. Visible spectra were measured and processed as follows. Spectra were first trimmed to wavelengths of 300nm-700nm (the visible light spectra) and smoothed using the PAVO package in R (with a smoothing span of 0.2 to maintain biological variation in reflectance measurements but reducing excess noise). Three colorimetric variables were calculated following previous work by other researchers in this area (Morehouse and Rutowski 2010a, Tigreros 2013, Espeset et al. 2019). In summary, midpoint (R50, associated with brightness) was the average percent reflectance between 300-375 and 450-550nm; LR50, associated with hue, was calculated as the associated wavelength to R50; and BR50, associated with saturation, was calculated by finding the slope of the line tangent to the point at R50. These measurements have been shown to be associated with pterin amount through comparisons of pterin-extracted wings and non-pterin extracted wings, and through comparisons of granule counts in scanning electron microscope images (Morehouse and Rutowski 2010b). Wings were photographed and measured from the basal tip to apex tip, and wing length was used as a proxy for body size.

### Dissections

Abdomens were added to approximately 1 mL 10% PBS buffer (Fischer Scientific). Testes and accessory glands were removed from males and spermatophores were removed from females under a dissecting scope in 1X PBS buffer. Reproductive organs were again stored in 10% PBS buffer and at −20°C until processed. Testes were weighed to the nearest microgram on a Mettler Toledo microbalance. Spermatophore counts were taken at time of dissection.

### Analyses

All analyses were conducted using R v.3.5.1. Linear models with population as a categorical factor were used on testes weight to investigate differences among populations, with post-hoc comparisons calculated using Tukey’s honest significant different (HSD) test. A similar set of analyses were used on spermatophore counts to examine population differences. Levene’s tests for homogeneity of variance were run as pairwise comparisons to determine differences in variance among sites in color variables for male wings (with respect to hypotheses described above and in Fig. 1).

Additional linear models were used to determine if colorimetric variables could predict testes weight. Because colorimetric variables were highly correlated amongst themselves, we ran three separate linear models for testes weight with each predictor colorimetric variable (R50, LR50, and BR50); each model included population as a categorical predictor and wing length as a covariate. We investigated the possibility that the relationship between color variables and testes size varied among populations by testing for an interaction between population and color variables. For all models, the distribution of residuals was examined for normality, and variance inflation factors were evaluated to ensure independence of predictors. Slopes and associated standard errors (from testes size predicted by color variables) were used to visualize differences among populations in the strength of the putative honest signal.

## Results

We caught and processed a total of 367 individuals from four different sites, two from northern Nevada (on the edge of the city of Reno, and Washoe Lake) and two from north-central California (the Davis Community Gardens, and Sutter National Wildlife Refuge), with numbers of individuals as follows: 59 males and 28 females from Reno, 68 males and 30 females from Washoe, 62 males and 27 females from Davis, and 54 males and 43 females from Sutter.

### Individual level differences

Brighter (R50), longer wavelengths (hue; LR50), and more saturated (BR50) wings were associated with larger testes sizes. In Fig. 2, we show relationships between testes weight and color variables separately for each site as well as across all sites and individuals (the solid black lines in Fig. 2), with results from the larger scale (across all individuals) as follows: R50, F_1,227_ = 8.06, R^2^ = 0.034, p = 0.005; LR50, F_1,227_ = 15.03, R^2^ = 0.062, p < 0.001; BR50, F_1,227_ = 4.58, R^2^ = 0.020, p = 0.033). Full models with site and wing length as additional variables (see Table 1) explained considerably more variation in testes size (compared to color variables alone, as in Fig. 2), but the most powerful variable in those models was wing length. We did not detect an interaction between color variables and collection site (R50, F_3,176_ = 0.21, p = 0.89; LR50, F_3,176_ = 0.81, p = 0.49; BR50, F_3,176_ = 0.27, p = 0.85), even though (as in Fig. 2) the association between color and testes size did appear to vary in magnitude among sites.

**Fig. 2.**
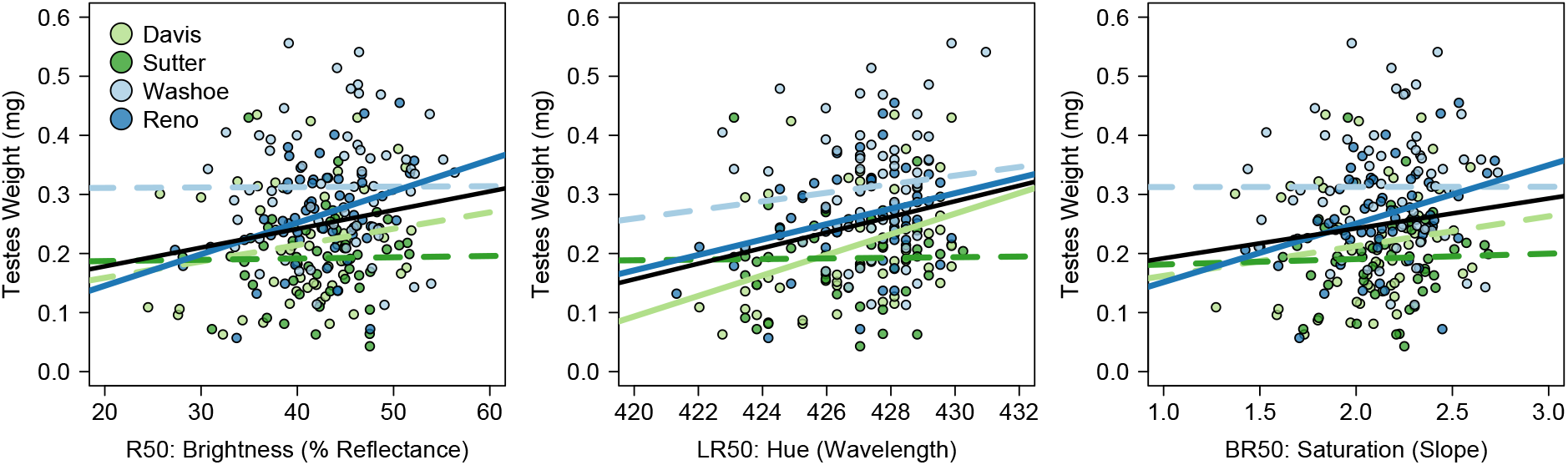
Teste weights by colorimetric variables (i.e. R50: midpoint; LR50: wavelength; and BR50: slope) for all male individuals from Davis, Sutter, Washoe, and Reno sites (see Fig. 1 map). Relationships across all individuals in each plot are represented by the solid black lines. Relationships for subsets of individuals (by site) are shown as colored lines, with relationships associated with *P* < 0.05 shown as solid lines. Slopes for the solid lines (across locations) are as follows, with standard errors in parentheses: R50, 0.0032 (0.0011); LR50, 0.013 (0.0034); BR50, 0.051 (0.024).

**Table 1.**
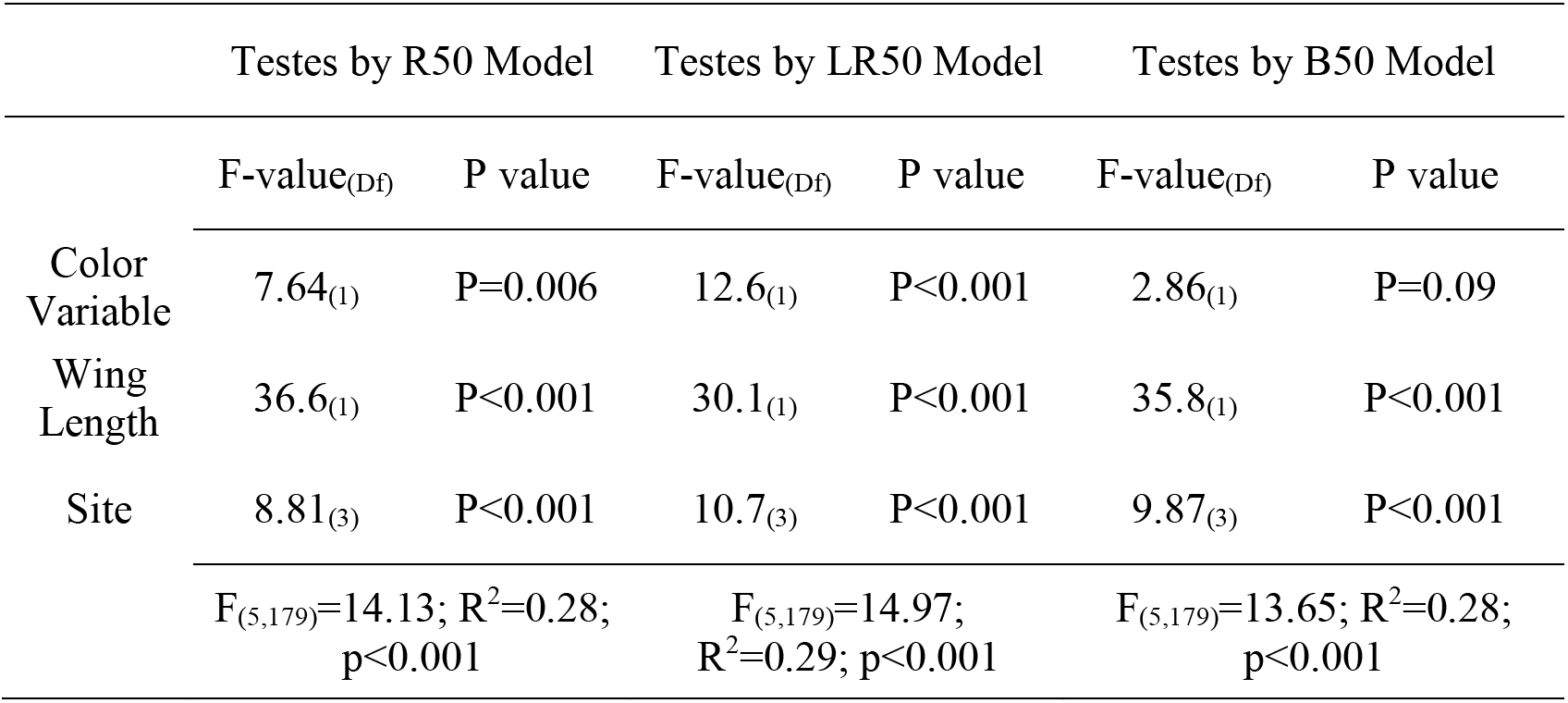
Results from linear models predicting testes size with color variables, wing length, and site as a categorical factor (see Fig. 1 for locations studied).

### Population level differences

Testes size varied across sites: the California sites (with greater proximity to agriculture) had overall smaller testes (and were similar to each other), while individuals from the Nevada sites had larger testes, with individuals from Washoe being the largest (Fig. 3A). In general, the Nevada sites are surrounded by less agriculture than the California locations, and the Washoe site was the least agriculturally-developed of all (specimens were collected near a wetland and terminal desert lake).

**Fig. 3.**
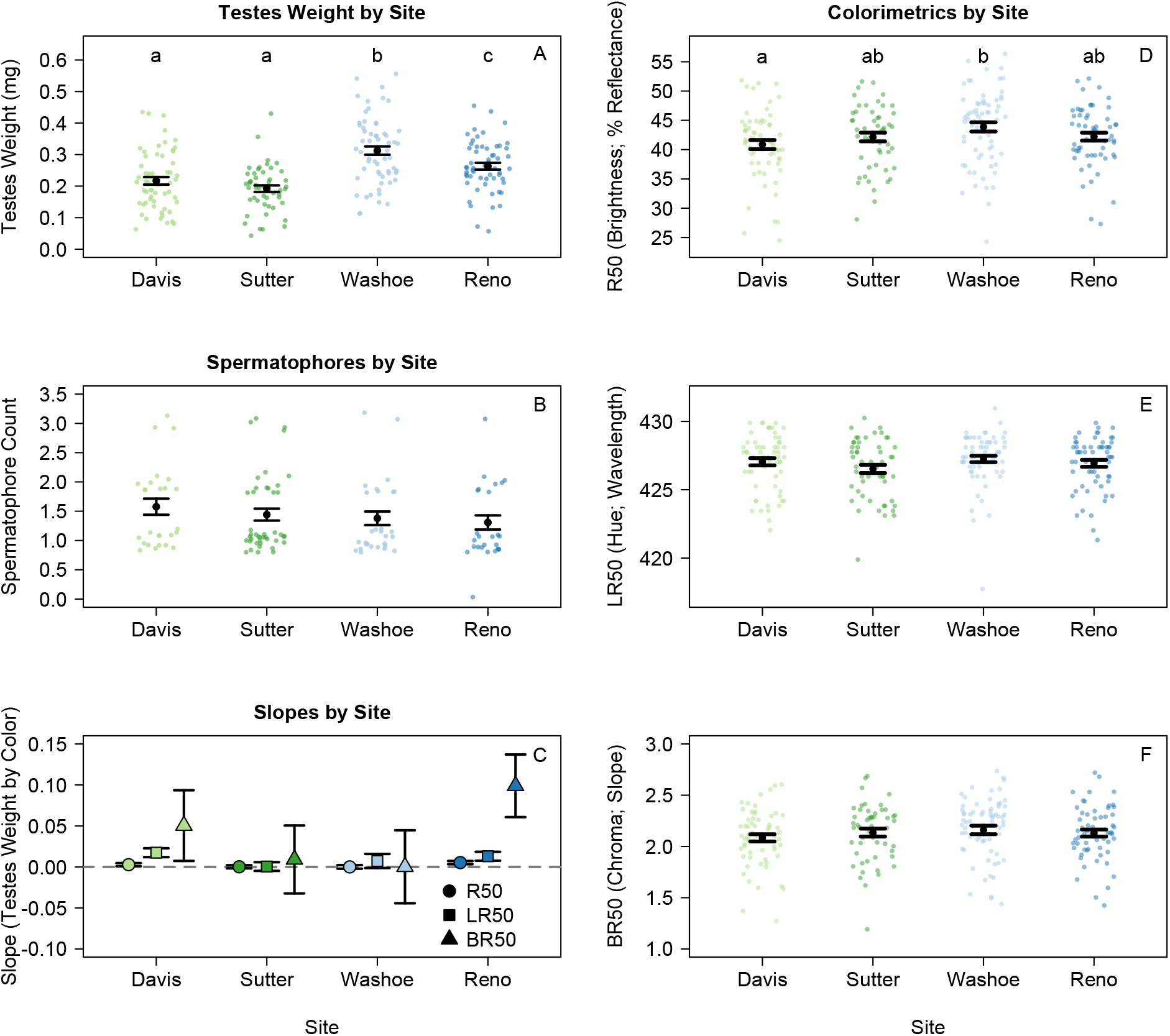
Within and among population variation in key variables studied, including reproductive traits (A and B) and colorimetrics (D-F). In panels A-E and F, points are jittered for visualization and lower case letters within plots are shown if post hoc comparisons indicated differences among populations at *P* < 0.05. Panel C shows among-population differences in the slope of testes weight by color variables (see legend for symbols) where more positive values suggest an honest signal (a relationship between testes weight and wing color).

All populations had at least one individual with three spermatophores and all specimens had at least one spermatophore found in dissections (except for one single, apparently unmated individual from our Reno site), with a median of one spermatophore dissected per population. Sites did not differ in the average number of spermatophores dissected from females (Fig. 3B) suggesting that populations at the different sites are similar in behavior with respect to the number of times females successfully mate.

As an alternative visualization of relationships between testes and color variables, slopes for each site from our above analyses of testes weight by R50, LR50, and BR50 variables (Fig. 2) are shown in Fig. 3C along with standard errors around the slope estimates. We found considerable variation in slopes among populations, with variables that differ from the zero line (i.e. standard errors not overlapping zero) suggesting situations in which wing color could be an indicator of the male reproductive trait. This was evident in the Reno and Davis populations (for BR50) in particular, and also for LR50.

We found variation among populations for all three color variables investigated, and that was particularly true for R50 (see independent contrasts in Fig. 3D), with the brightest individuals being from our Washoe site, Davis having the least bright individuals, and our other two sites having intermediate brightness (Fig. 3D). We did not detect differences among sites in hue or saturation in the simple contrasts used for visualization in Fig 3 (panels E and F), but it should be noted that an overall effect of site on those color variables was detected in full models (with other covariates) reported in Table 1. Using Levene’s test for homogeneity of variance, we did not detect differences among locations in the three wing color variables.

## Discussion

Evidence that humans are affecting mating signals and mating systems through nutritional changes to natural and modified landscapes is growing (Snell-Rood et al. 2015, Goos et al. 2016, Espeset et al. 2019). In this study we looked at differences in the relationship between wing coloration and testes size among populations in the cabbage white butterfly, a species with extensive exposure to anthropogenic habitats. We collected specimens from two sites associated with low cropland and two sites associated with high cropland to both quantify the extent to which wing color could be an indicator of a reproductive trait in this system and to ask if any relationship between color and testes size might vary among locations. The present study (with four locations sampled) was not designed to explicitly test for landscape-level factors (e.g. climactic variation, host-plant availability, etc.) affecting reproductive traits, rather our goal was to open a window onto the potential for evolutionarily-important, among-population differences.

Honest signaling in secondary sexual traits in the cabbage white butterfly has been previously suggested (Morehouse and Rutowski 2010b), although in that study male quality was not quantified. In another study (Tigreros 2013), a relationship between spermatophore content and coloration was not detected, however those authors did not measure color and reproductive tissue from the same individual. Our results contribute to this area of investigation by providing evidence that a reproductive trait (testes size) is indeed positively associated with coloration when measurements are taken from the same individuals. This could indicate that coloration in cabbage white butterflies is, in fact, an honest indicator of reproductive investment, although that relationship (between color and reproductive investment) explains relatively little variation (Fig. 2).

When considering site-level relationships between color and testes, we did not detect an overall interaction between site and color variables (predicting testes size), but we did find that the color-testes relationship was notably stronger in some locations (see slopes in Fig. 3C, and site-specific lines in Fig. 2). Reno had the strongest association of testes size with each color variable, and Davis had a positive trend, while Washoe and Sutter sites showed much weaker or nonexistent relationships between wing coloration and testes size (Fig. 2, Fig. 3C). In terms of proximity to agriculture and potential anthropogenic nutrient inputs, Sutter is our most intensely modified site, followed by Davis, then Reno and finally Washoe with the least agricultural influence. Given the data currently in hand, it is not clear why our two intermediate sites (neither most intensive crop areas nor the areas with the least cropland) showed the strongest relationships between wing color and a reproductive trait. We discuss possible interpretations below in the spirit of raising hypotheses for future study.

The difference in the relations between wing color and testes size among sites could suggest that individuals from the very high agricultural land-use area (Sutter) have been exposed to relatively high amounts of nitrogen for a longer period of time than any other site, relaxing the value of investing in an expensive, sexually-selected secondary trait that no longer acts as an honest signal of mate quality. Our Davis site may still hold some trait honesty but less than our Reno site, which has been exposed to agricultural land and fertilizer deposition, but possibly for a shorter amount of time. The fact that our site with the lowest assumed nitrogen availability (Washoe) did not show trait honesty is perplexing and highlights the need for further investigation and direct measurement of levels of available nitrogen in the landscape. The Washoe site was low density (in terms of adult butterflies), thus another possibility might be investment in testes rather than wing color in an area were competition for mates is low.

Finally, it is worth noting that further complexity might derive from the fact that the nitrogen-flooded areas could also have higher nitrogen-containing nectar sources (i.e. higher concentrations of amino-acids (Baker and Baker 1977, Gardener and Gillman 2001)), with which individuals can replenish nutrient depletions as adults and can increase reproductive success or fecundity (Jervis and Boggs 2005, Mevi-Schütz and Erhardt 2005). More studies are needed to fully understand the dynamics of increased nitrogen, including availability and sequestration at different life stages associated with populations in different habitats (Bertazzini and Forlani 2016).

Additional studies are also needed on relationships between colorimetric variables in the visual spectra and female vision, which are not completely understood in butterflies (Stavenga and Arikawa 2006); however, we do know that females should be able to differentiate individuals that vary in color within the visible range, which is what we have focused on (Morehouse and Rutowski 2010b). We found positive relationships between testes size and color, suggesting that it would indeed be beneficial for females to differentiate between individuals of high and low quality based on color. A behavioral analysis of female mate selection across populations in areas with differing nitrogen availability would help us understand these signals better.

In our study, we focused on habitats that potentially differ in nutrient availability, but it is important to remember that there are numerous other factors that could be causing the differences in testes size we see in this study. Due to the labor-intensive nature of quantifying both mate quality (testes size) and wing color, our investigations here were limited to four locations. Thus we infer that the relationship between mate quality and the mating signal likely varies among locations, but we cannot yet explain that variation, and expect that it involves many factor not measured here, e.g. pesticide use, climate, elevation, maternal effects, and others. The importance of additional, unmeasured factors is also supported by the observation that individuals from agricultural areas tended to have smaller testes and less colorful wings (Fig. 3). The possibility of multiple factors driving variation in reproductive traits combined with the possibility that different populations are at different points along an evolutionary trajectory (possibly leading to the de-valuing of an honest signal) highlights the complexity and value of studying sexually selected signals in the age of the Anthropocene.

## Acknowledgments

This material is based upon work supported by the National Science Foundation Graduate Research Fellowship under Grant No. 1447692 to AE. Thank you to Dr. Emilie Snell-Rood and her lab for space and microscope use for reproductive dissections, thanks for Dr. Anne Leonard for use of her JAZ spectrophotometer, and thanks to Dr. Art Shapiro for advice on when and where to find the most butterflies. We are grateful to lab technicians Taylor Bradford and Dominic Zullo, as well as for thoughtful comments from Joaquin Baixeras and an anonymous reviewer.

